# Microbial aromatic amino acid metabolism is modifiable in fermented food matrices to promote bioactivity

**DOI:** 10.1101/2023.12.21.572869

**Authors:** Mikaela C. Kasperek, Adriana Velasquez Galeas, Maria Elisa Caetano-Silva, Zifan Xie, Alexander V. Ulanov, Michael R. La Frano, Suzanne Devkota, Michael Miller, Jacob M. Allen

## Abstract

Ingestion of fermented foods impacts human immune function, yet the bioactive food components underlying these effects are not understood. Here, we interrogated whether fermented food bioactivity could be traced to a class of microbial metabolites derived from aromatic amino acids (ArAA), termed aryl-lactates. Using targeted metabolomics, we established that the aryl-lactates phenyllactic acid (PLA), 4-hydroxyphenyllactic acid (4-HPLA), and indole-3-lactic acid (ILA), are present in varying concentrations across a wide range of commercially available fermented foods, including many vegetable and dairy ferments. After pinpointing fermented food-associated lactic acid bacteria (LAB) that produce high levels of aryl-lactates (e.g., *Lactiplantibacillus plantarum*), we utilized our knowledge of LAB metabolism to identify fermentation conditions (added cultures [e.g., *L. plantarum*] and metabolic co-factors [e.g., aryl-pyruvates]) to increase aryl-lactate production in food matrices up to 5×10^3^ fold vs. standard fermentation conditions. Next, using *ex vivo* reporter assays, we found that a variety of food matrix conditions optimized for aryl-lactate production exhibited enhanced agonist activity for the human aryl-hydrocarbon receptor (AhR) as compared to standard fermentation conditions and/or commercial brands. Moreover, we determined that strategies to enhance aryl-lactates effectively maintained food matrix AhR bioactivity across 4 weeks of storage. Reduced microbial-induced AhR activity has emerged as a hallmark of many chronic inflammatory diseases, thus we envision strategies to enhance microbially produced aryl-lactates and thus AhR bioactivity of fermented foods can be leveraged to improve human health.

## INTRODUCTION

The process of fermenting foods has traditionally been utilized to extend shelf life of perishable items, but more recently has been investigated for its proposed health-promoting effects upon consumption^1, 2^. Fermented foods contain desired microbial growth and enzymatic conversions resulting in transformation of food components^3^. Various fermented foods are commercially available including dairy ferments such as yogurt and kefir, vegetable ferments including sauerkraut and kimchi, fermented tea (kombucha), and even fermented meats like salami. Fermentation involves either initiation with starter culture or independent (wild/spontaneous) fermentation, often involving lactic acid bacteria (LAB)^4^. LAB have long been recognized for their pivotal role in improving food safety and extending shelf life, but their significance extends well beyond preservation, including their suggested involvement in shaping the gut microbiota upon consumption, where they may confer a range of benefits^3, 5–10^. Recent work indicates that the consumption of a diet rich in LAB-containing fermented foods in otherwise healthy individuals can dampen circulating inflammatory cytokines^11^. However, neither the type of fermented food, fermentation microbe(s), nor the specific microbial metabolite(s) within the fermented foods that contribute to health-promoting effects have been uncovered.

Previous work indicates that LAB fermentation increases bioavailability of nutrients and enhances the synthesis of vitamins and biologically active compounds within the food matrix^3, 12^. Outside a small body of work, the sources of bioactivity within a fermented food matrix have not been thoroughly characterized. Recent studies have identified gut microbiome-derived LABs ability to metabolize aromatic amino acids into a variety of downstream metabolites with bioactive potential. Phenyllactic acid (PLA), indole-3-lactic acid (ILA), and 4-hydroxyphenyllactic acid (4HPLA); collectively termed as aryl-lactates in this paper, have emerged as critical metabolites derived from gut microbes that impact human physiology, including growth, development, and immune function^13^. PLA, 4HPLA and ILA are derived from aromatic amino acids (ArAA) phenylalanine (Phe), tyrosine (Tyr), and tryptophan (Trp), respectively, via enzymes aromatic aminotransferase (ArAT) and microbial phenyllactate dehydrogenase (fLDH)^13, 14^. Data from our laboratory and others have uncovered metabolic and immunomodulatory activity of aryl-lactates, including upregulation of serum concentrations of 4HPLA and ILA after exercise training likely from gut microbiota production, and attenuation of inflammation by ILA treatment in various settings^15–23^. While mechanisms are not clear, evidence suggests that aryl-lactates can bind immunomodulatory receptors including the human hydroxycarboxylic acid receptor 3 (HCAR3), and aryl hydrocarbon receptor (AhR)^24, 25^. While studies have identified specific LABs such as *L. plantarum* as aryl-lactate producers^26, 27^, there is limited evidence that these metabolites can be found within fermented foods. Moreover, no studies to our knowledge have compared the quantity of microbial aryl metabolites between food types or determined if their metabolism can be manipulated to augment the bioactivity of a whole food matrix^16, 28–30^.

This study first characterized concentrations of bioactive aryl-metabolites in a variety of commercially available fermented foods, identifying wide-ranging aryl-lactate concentrations across food types and brands. We then utilized our knowledge of microbial ArAA metabolism to modify production of aryl-lactates within monoculture and whole fermented food settings. We ultimately found that manipulating microbial ArAA metabolism towards higher aryl-lactate production significantly enhanced the bioactivity (AhR activity) of fermented food matrices, exemplifying immunomodulatory potential upon consumption of these foods.

## RESULTS

### Fermented foods identified as a source of microbial-derived aryl-lactates: PLA, 4HPLA, and ILA

Fermented foods were purchased from a local grocery (Champaign, IL) to include a variety of dairy (e.g., Greek yogurt, kefir), vegetable (e.g., Sauerkraut), and miscellaneous (e.g., animal protein, fish sauce) fermented products. We first examined whether aryl-lactates were present and their concentrations in these foods using a LC/MS/MS targeted metabolomics method (**Fig. 1A**, Fig. S1). We identified aryl-lactates derived from Phe (phenyllactic acid-PLA), Tyr (4-hydroxyphenyllactic acid-4HPLA) and Trp (indole-3-lactic acid-ILA) to be present in nearly all fermented foods tested, but in varying amounts depending on the food type and source (**Fig. 1B-E**). The dairy ferments with the highest aggregate levels of aryl-lactates (PLA+4HPLA+ILA) included kefir and select Greek yogurts (**Fig. 1B**). The vegetable ferments with the highest aggregate aryl-lactates were fermented carrots and select brands of sauerkraut (**Fig. 1B**). Miscellaneous ferments (e.g., fish sauce, kombucha) contained on average ∼4-6-fold lower aryl-lactates compared to vegetable or dairy ferments (**Fig. 1B**). PLA (0.17-28.4 µg/mL; **Fig. 1C**) and 4HPLA (0.4-43.7 µg/mL; **Fig. 1D**) were the most representative aryl-lactates across all fermented food sources. ILA (0-2.8 µg/mL; **Fig. 1E**) was the least present aryl-lactate in all fermented foods, representing a bias towards Phe and Tyr metabolism in whole food matrices or related to the concentrations of the ArAAs in the whole foods (**Fig. 1F**, Fig S2). While the total aryl-lactate concentration in dairy vs vegetable ferments were not different, normalized concentrations of aryl-lactates were dependent on the source of the whole food. Dairy ferments exhibited significantly more 4HPLA, while vegetable ferments contained more PLA (**Fig. 1F**; p<0.001). We also determined aryl-lactate concentrations per serving size.^31^ Because fermented dairy serving sizes are larger than those of fermented vegetable products, an average serving of dairy ferments contains more aryl-lactates than one serving of vegetable ferments (**Fig. 1G**; p<0.01).

**Figure 1.**
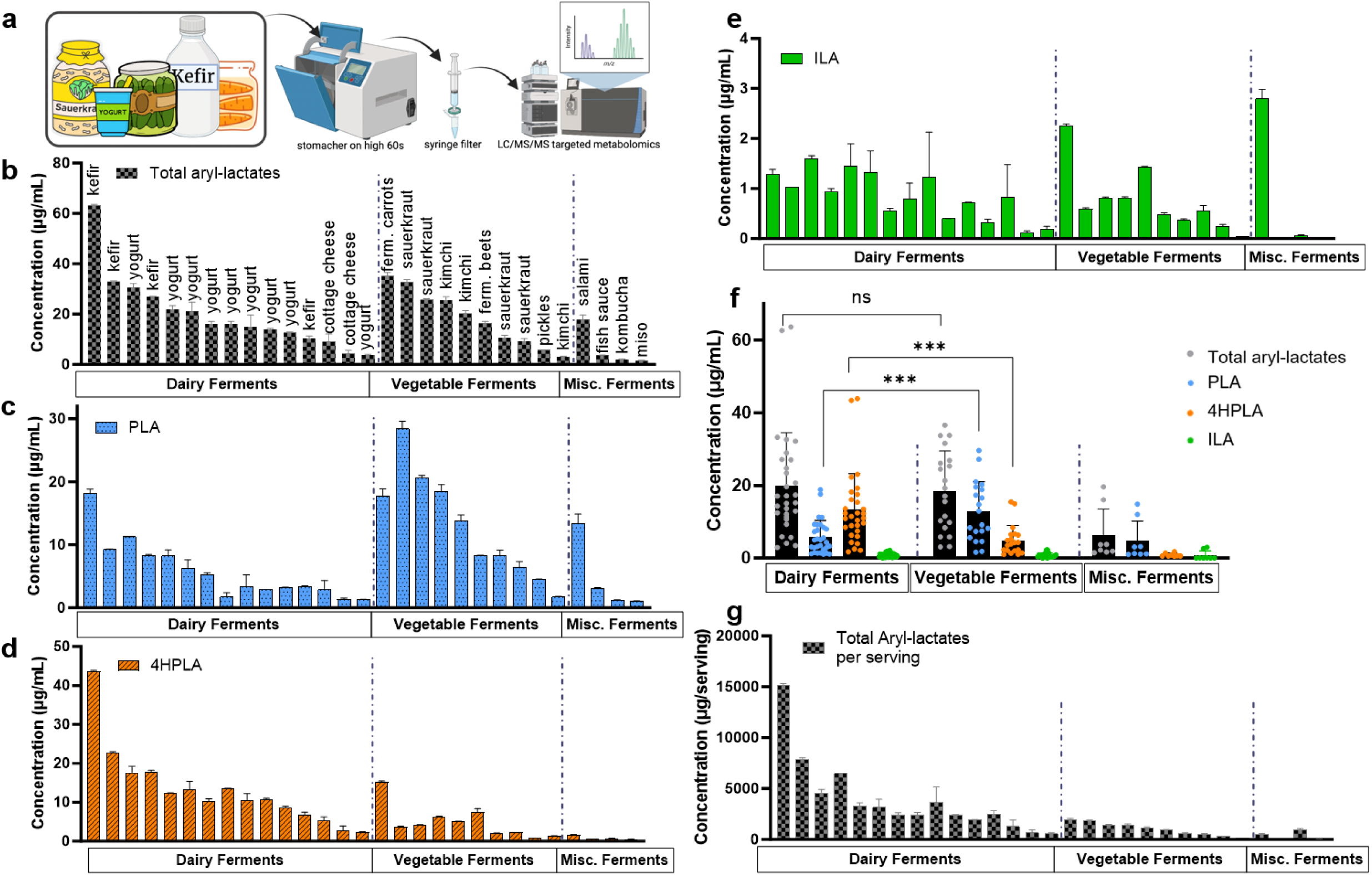
Fermented foods are a source of microbial-derived aryl-lactates. (**a**) Commercially available foods were homogenized to elute metabolites in a stomacher for 60s, syringe filtered, and analyzed via LC/MS/MS metabolomics using aryl-lactate standards. (**b**) total aryl-lactates (PLA+4HPLA+ILA) (**c**) Phenyllactic acid (PLA), (**d**) 4-hydroxyphenyllactic acid (4-HPLA), and (**e**) indole-3-lactic acid (ILA) concentrations are shown in µg/mL. Foods are ordered by descending total aryl-lactate concentration within fermented food categories (dairy, vegetable, miscellaneous) with fermented food type showed, B-E,G have all foods in the same order. (**f**) Commercially available fermented foods separated and combined according to type of ferment (fermented dairy products, left; fermented vegetables, middle; miscellaneous ferments, right). Total aryl-lactates is the sum of PLA, 4HPLA, and ILA for each fermented food. Statistics using 1-way ANOVA showing comparison between dairy and vegetable ferments total aryl-lactates, PLA, and 4HPLA ***p<0.001. (**g**) Total aryl-lactates (PLA+4HPLA+ILA) per serving of commercially available fermented foods. 1 serving considered 8 oz kefir and kombucha, 6 oz yogurt, 1/4c vegetable ferments, 1Tbsp fish sauce, and 2 tsp miso paste. Statistics using unpaired T-test comparing average aryl-lactates in 1 serving vegetable vs. dairy ferments **p<0.01. All data displayed as mean +/- SEM.

### Bacteria strains with aryl-lactate producing capacity are present in commercially sold fermented food products

After identifying the presence of PLA, 4HPLA, and ILA in various fermented foods, we next investigated the potential bacterial sources of aryl-lactate production within fermented foods. We cultured the ferments anaerobically and isolated DNA from bacteria isolates. Using qPCR, we targeted a panel of lactic acid bacteria (LAB) species commonly found in fermented foods.^32^ qPCR identified a variety of LABs across food sources. Within the foods with the highest aryl-lactates, the most common and abundant bacteria isolates were *Lactiplantibacillus plantarum* (*L. plantarum*), followed by *Lactobacillus brevis* and *Lacticaseibacillus paracasei* (Fig S3). We next sought to determine ability of select LAB strains (including 7 strains of *L. plantarum*) found in fermented foods to directly produce aryl-lactates. After 24hr monoculture in standard MRS or GM17 media, supernatant was analyzed using LC/MS/MS for concentrations of aryl-lactates (**Fig. 2A-E**). In accordance with its presence in foods with high amounts of aryl-lactates, all tested *L. plantarum* strains produced significantly higher amounts of total aryl-lactates, PLA, and ILA compared to other LABs tested (**Fig. 2B-C, E** p<0.0001).

**Figure 2.**
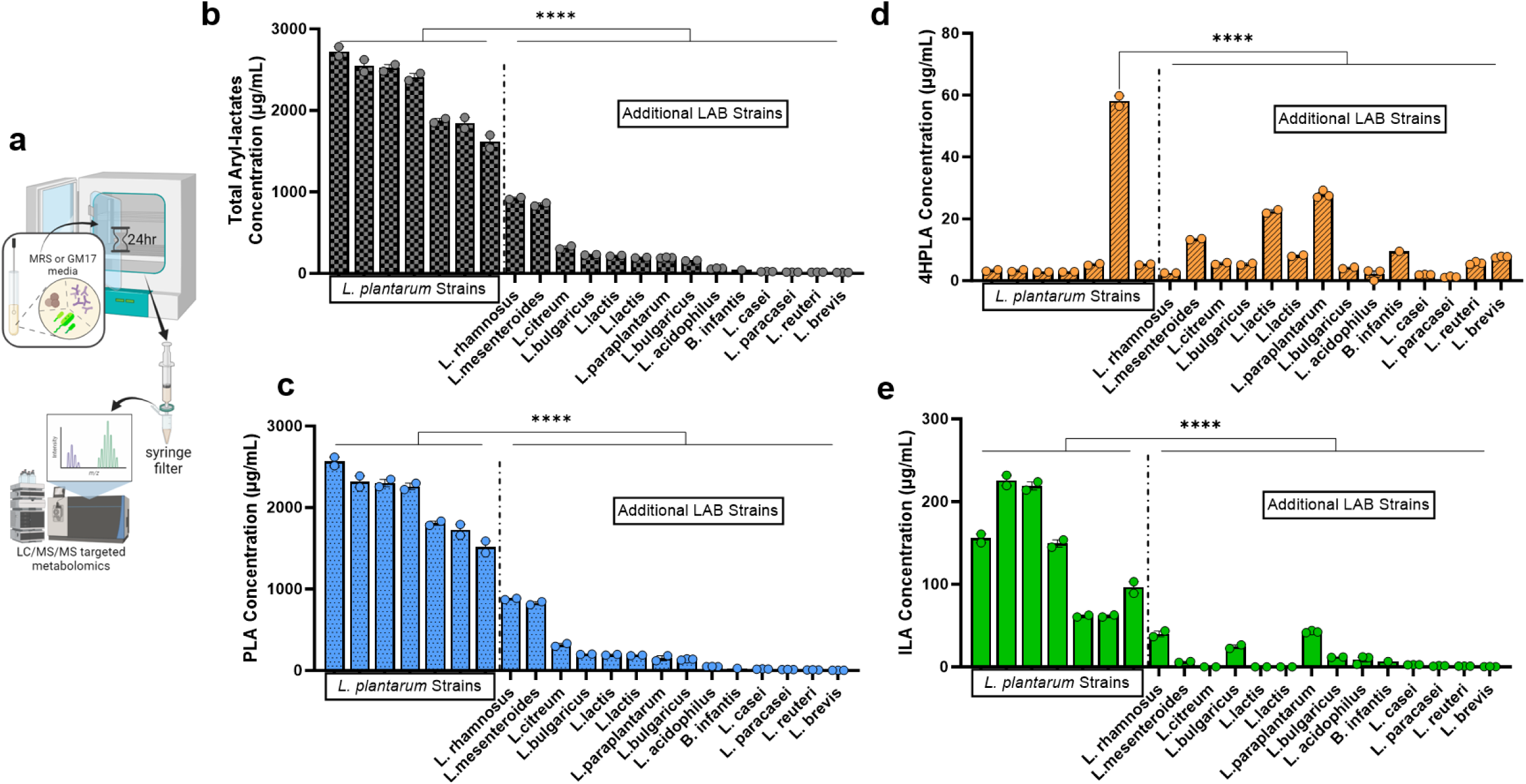
Select lactic acid bacteria (LAB) found in commercially available fermented foods produce aryl-lactate metabolites in monoculture. (**a**) Supernatant from bacteria monoculture were syringe filtered and analyzed via LC/MS/MS targeted metabolomics using aryl-lactate analytical standards. (**b**) Total aryl-lactates, (**c**) Phenyllactic acid (PLA), (**d**) 4-hydroxyphenyllactic acid (4-HPLA), and (**e**) Indole-3-lactic acid (ILA) concentrations (µg/mL) in select lactic acid bacteria monocultures analyzed via LC/MS. All strains ordered from greatest to least concentration of total aryl-lactates. 7 *Lactiplantibacillus plantarum* (*L. plantarum*) strains (left) and additional LAB strains (right, *L. rhamnosus*, *Leuconostoc mesenteroides*, *Leuconostoc citreum*, *L. bulgaricus*, Lactococcus lactis, L. paraplantarum, an additional *L. bulgaricus* strain, *L. acidophilus, Bifidobacterium infantis, L. casei, L. paracasei, L. reuteri,* and *L. brevis*). Statistical analysis performed using 1-way ANOVA with Dunnett’s multiple comparison’s test with each *L. plantarum* strain compared to additional LAB strains. Only significant values for all additional LAB strains shown at p<0.05. ****p<0.0001. Data represented in mean +/- SEM.

### Select co-factor and precursor metabolites shift LAB metabolism towards increased production of aryl-lactates

We next worked to optimize the metabolism of ArAA to favor the production of aryl-lactates in monoculture. We focused on the LAB *L. plantarum* because of its capability to produce high concentrations of aryl-lactates in monoculture as well as its commonality within both wild and non-wild fermented foods.^33^ We also chose to investigate the LAB *B. infantis* not only to determine if different bacteria react in varying manners to metabolism manipulation, but also because of its prevalence and physiological importance in the human gut microbiome.^34, 35^ Since alpha-ketoglutarate (AKG) is a known nitrogen acceptor in the metabolism of amino acids (**Fig. 3A**) as well as citrate (CIT) potentially act as a stimulator of ArAA catabolism,^36^ we added both cofactors individually and in combination to *L. plantarum* and *B. infantis* monoculture to determine if they increase the production of aryl-lactates and could potentially be limiting factors in the metabolism of these metabolites. The combination of both AKG and CIT increased the production of aryl-lactates in *L. plantarum* monoculture by 42-86%, with PLA exhibiting the most robust increase compared to monoculture with no additives (**Fig. 3B**). Aryl-lactates production by *B. infantis* were significantly more sensitive to AKG and CIT compared to *L. plantarum*, exhibited by >7,000-fold increase in PLA and >900-fold increase in ILA compared to the control *B. infantis* culture with no AKG or CIT (**Fig. 3C**).

**Figure 3.**
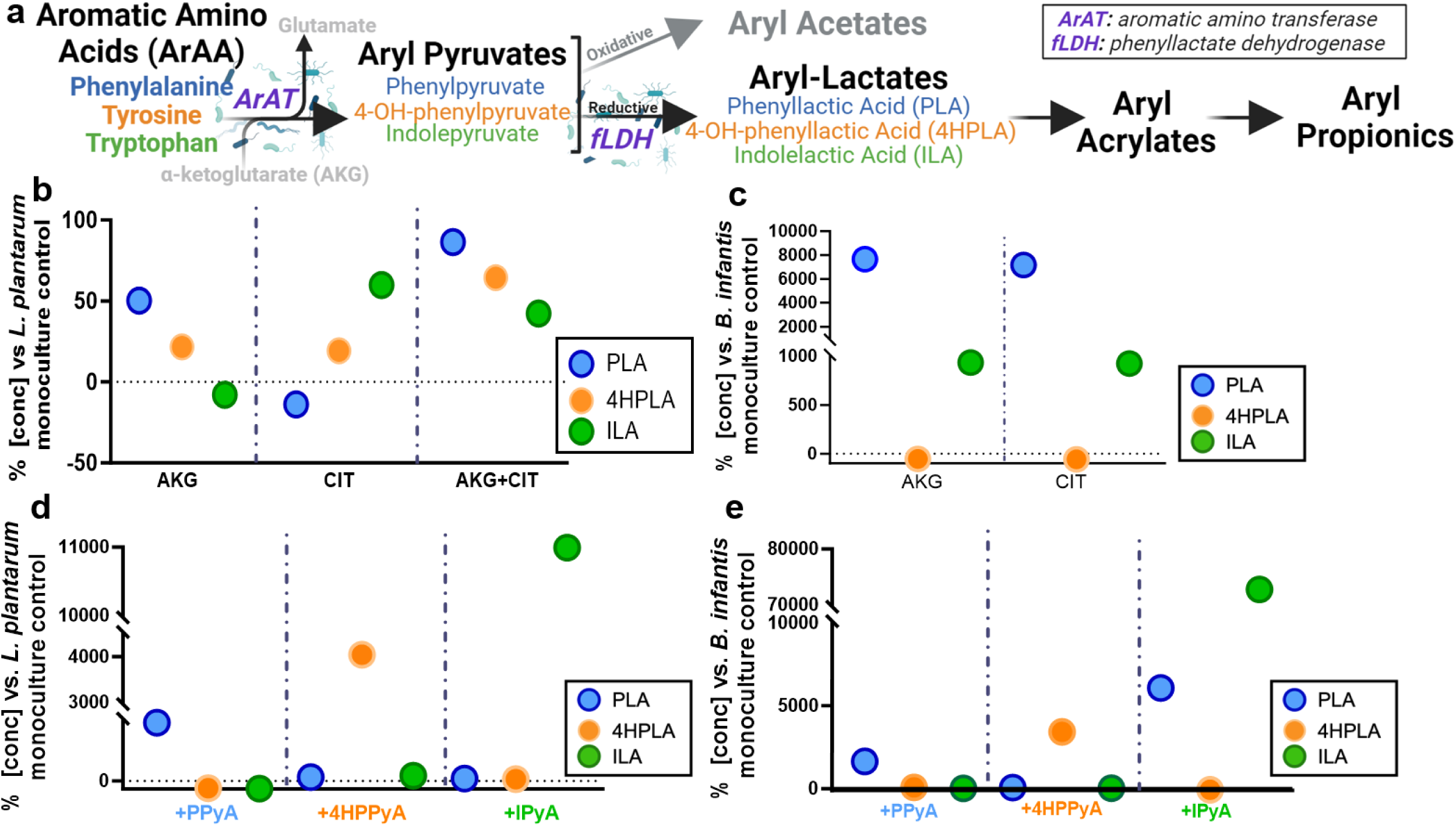
Addition of co-metabolites to Lactic acid bacteria (LAB) monocultures promotes the production of aryl-lactates. (**a**) Aromatic amino acid (ArAA) microbial metabolism pathway showing aromatic amino acid metabolism via aromatic amino transferase (ArAT) to aryl-pyruvates, then aryl-pyruvates are metabolized via microbial enzyme phenyllactate dehydrogenase (fLDH) to aryl-lactates and more downstream to additional aryl metabolites. (**b**) *Lactiplantibacillus plantarum* (*L. plantarum*) or (**c**) *Bifidobacterium longum spp. infantis* (*B. infantis*) monoculture inoculated with alpha-ketoglutarate (**AKG**) and/or trisodium citrate dehydrate (**CIT**) and syringe filtered supernatant analyzed after 24hr via LC/MS/MS. (**d**) *L. plantarum* or (**e**) *B. infantis* monoculture inoculated with phenylpyruvic acid (**PPyA**), 4-hydroxyphenylpyruvic acid (**4HPPyA**), or indole-3-pyruvic acid (**IPyA**) for 24 hours prior to LC/MS/MS targeted analysis of syringe filtered supernatant. All data shown as % concentration of aryl-lactates in treatment condition compared to respective monoculture with no additives.

ArAAs are metabolized by select bacteria to aryl-pyruvates via the aromatic aminotransferase (ArAT) enzyme and are subsequently metabolized to aryl-lactates via microbial enzyme phenyllactate dehydrogenase (fLDH) (**Fig. 3A**). In an effort to enhance the production of aryl-lactates in LAB cultures, we next added aryl-pyruvates (phenylpyruvate [PPyA], 4-hydroxyphenylpyruvate [HPPyA], or indole-3-pyruvate [IPyA]) to *L. plantarum* and *B. infantis*. Interestingly, in almost all cases when the upstream aryl-pyruvate was added, the respective downstream aryl-lactate was robustly increased while the other two aryl-lactates exhibited slight, or no change compared to control culture (**Fig. 3D-E**). For example, when indole pyruvate (IPyA) was added to either a *L. plantarum* or *B. infantis* monoculture, ILA increased in production ∼10,000% and ∼7,500%, respectively, compared to *L. plantarum or B. infantis* monoculture without indole-pyruvate addition (**Fig. 3D-E**).

### Manipulation of ArAA metabolism transfers to whole fermented food matrices and is optimized further across commercially relevant storage times

We next examined whether aryl-lactate metabolism could be modified in a whole fermented food matrix. We first focused on yogurt because 1) we established it as a source of aryl-lactates and 2) it has a known and well-established starter cultures (*S. thermophilus*, *L. bulgaricus*) compared to other dairy and vegetable ferments. Yogurt (*Starter culture only: 0.4% w/v of 1×10*^7^ *CFU/mg L. bulgaricus, S. thermophilus*) was fermented for 5 hours prior to sampling at three timepoints: immediate post-5hr fermentation, 1-week cold storage and 4-weeks cold storage (**Fig. 4A**). Aryl-lactate concentrations in starter culture only conditions were compared to yogurt conditions where we added bacterial strains, co-factors and metabolites to the starter culture: AKG and/or CIT, *B. infantis* or *L. plantarum* alone, individual aryl-pyruvates (PPyA or HPPyA or IPyA), pyruvate blend (PPyA+HPPyA+IPyA), and *L. plantarum* or *B. infantis* with pyruvate blend. First, we examined aryl-lactate production immediately post 5hr fermentation (**Fig. 4B**). We found that AKG or CIT marginally increased the production of aryl-lactates compared to control yogurt (4-27%). The addition of LABs *B. infantis* or *L. plantarum* robustly upregulated aryl-lactate production with 3.4-fold change on average compared to control yogurt. For example, we observed a 1,000% increase in PLA in yogurt with *L. plantarum* added (**Fig. 4B**). Mirroring what we observed in LAB monoculture, addition of individual aryl-pyruvates to yogurt robustly enhanced each respective downstream aryl-lactate (e.g., added IPyA: ∼25-fold increase in ILA), while the production of other aryl-lactates exhibited less change (e.g., added IPyA: 0.24-fold increase in PLA, 0.26-fold increase in 4HPLA) (**Fig. 4B**). Meanwhile, adding all aryl-pyruvates to yogurt culture together increased all downstream aryl-lactates on average 10-fold vs yogurt control where no aryl-pyruvates were added **(Fig. 4B**). The most effective conditions tested for promoting aryl-lactate concentrations in a whole food matrix was the addition of bacterial aryl-lactate producer (*L. plantarum* or *B. infantis*) in combination with a blend of aryl-pyruvates. The addition of *L. plantarum* and aryl-pyruvate blend to yogurt with only starter culture robustly increased all aryl-lactates [PLA (2,260%), 4HPLA (580%), and ILA (3,312%)] as compared to yogurt fermented with starter culture only (**Fig. 4B**, p≤0.0001).

**Figure 4.**
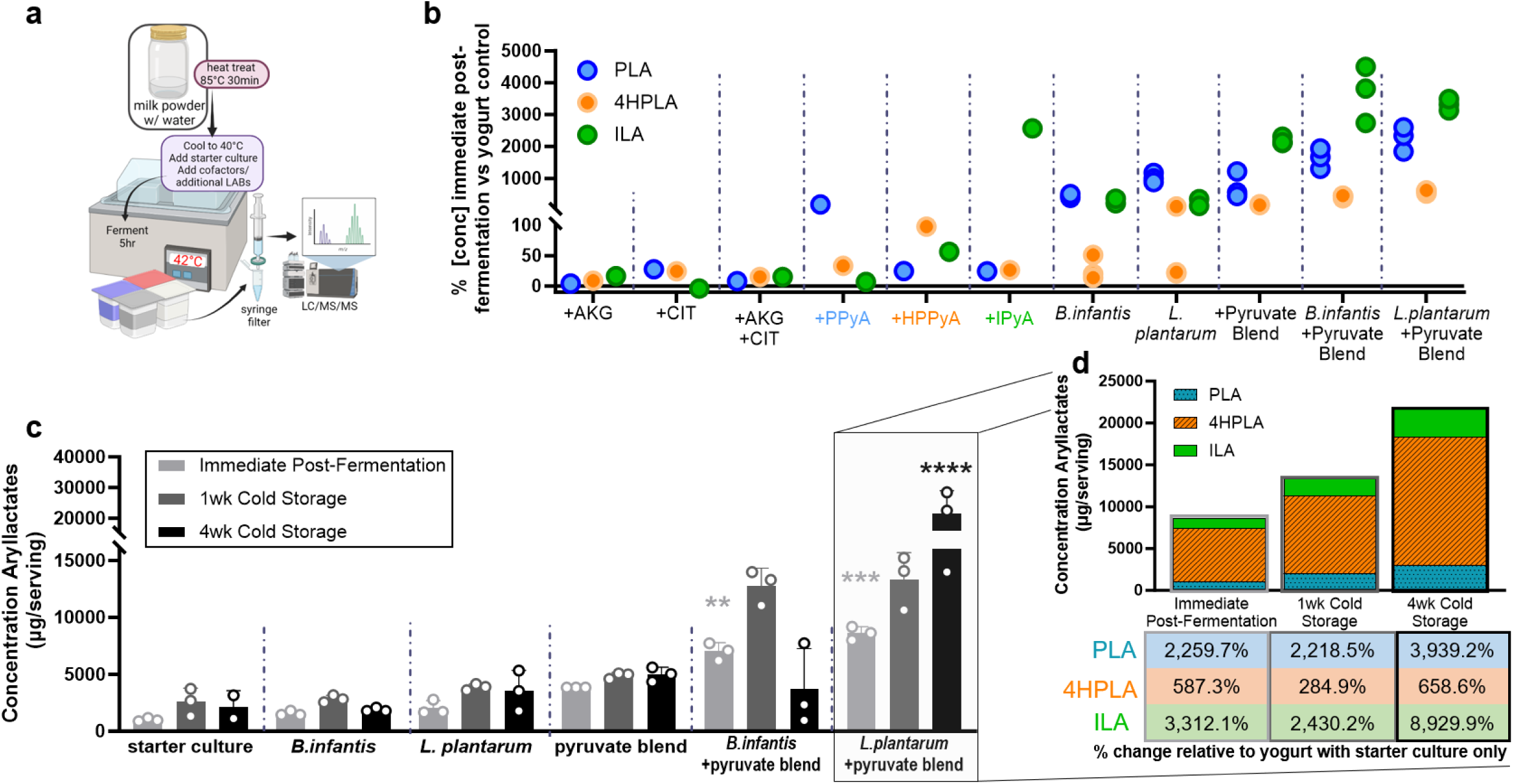
Aromatic amino acid metabolism is modifiable in a whole-food matrix to promote aryl-lactate production. (**a**) Yogurt was made by mixing milk powder and sterile water and heat treating the solution for 30min prior to being cooled to 40°C and starter culture (*Strep. Thermophilus*, *L. bulgaricus;* Chr-Hansen) plus applicable treatment were added before fermenting at 42°C for 5hr. All yogurt samples were syringe-filtered before analyzing via LC/MS/MS. (**b**) Yogurt with standard starter culture was fermented for 5hr with treatments including added alpha ketoglutarate (AKG) and/or citrate (CIT), aryl-pyruvates (phenylpyruvic acid (PPyA), 4-hydroxyphenylpyruvic acid (HPPyA), indole-3-pyruvic acid (IPyA)) and *B. infantis* or *L. plantarum* with and without blend of aryl-pyruvates (pyruvate blend; PPyA+HPPyA+IPyA). Phenyllactic acid (PLA), 4-hydroxyphenyllactic acid (4HPLA), and indole-3-lactic acid (ILA) percent concentration vs. yogurt control (standard starter culture only) is shown. (**c**) Concentration of total aryl-lactates per 6oz serving of yogurt via metabolomics analysis shown with various treatment groups and at three points over time: immediate post-5hr fermentation, fermentation and 1wk cold storage, and fermentation and 4wk cold storage. Statistical analysis performed using 2-way ANOVA with Dunnett’s multiple comparisons test with all treatments compared to yogurt with starter culture at same timepoint. Only significant values shown at p<0.05. **p<0.01, ***p<0.001, ****p<0.0001 (**d**) Concentration of aryl-lactates stacked to show ratio of PLA, 4HPLA, and ILA in yogurt serving across time points in yogurt with *L. plantarum* and pyruvate blend. Below each time point, percent change relative to yogurt with starter culture only is listed. Data displayed as mean +/- SEM.

Storage conditions and time can impact bioactive metabolites in food.^37^ Thus, we next tested: 1) whether total aryl-lactates (per 6oz serving of yogurt) changed over a commercially relevant storage time (1-4 weeks) and 2) if conditions best optimized to produce aryl-lactates immediately after fermentation could improve aryl-lactate maintenance across storage time (**Fig. 4C-D**, Fig S4A-B). In the starter culture only condition, aryl-lactates were maintained at similar levels after 4 weeks of storage compared to immediately post fermentation, indicating that aryl-lactate concentrations are relatively stable during 4wks cold storage. The addition of select aryl-lactate producing bacteria (*B. infantis, L. plantarum*) did not significantly increase the production of aryl-lactates, nor did it impact total aryl-lactate concentrations across cold storage (**Fig. 4C**; p>0.05). Similarly, the addition of pyruvate blend alone did not change total aryl-lactate concentration compared to yogurt with starter culture only. The production of aryl-lactates within yogurt significantly increased when an aryl-lactate producing bacteria was added in conjunction with aryl-pyruvate blend immediate post-fermentation period (**Fig. 4C**; *B. infantis* + pyruvate blend p<0.01, *L. plantarum* + pyruvate blend p<0.001). *L. plantarum* added in combination with the aryl-pyruvate blend was found to be most effective at maintaining aryl-lactates during cold storage. For example, *L. plantarum* + aryl-pyruvates yielded ∼9,000% fold increase in ILA after 4 weeks of cold storage compared to the starter culture only (**Fig. 4D**).

Next, we tested if aryl-lactate production could be fostered in a different fermented food matrix, sauerkraut. Unlike yogurt, sauerkraut is produced from a vegetable (cabbage) and undergoes fermentation from resident bacteria found naturally on the food (i.e., no starter cultures are added). To test this, we added a variety of metabolic co-factors (e.g. AKG and/or CIT, aryl-pyruvates) with or without the addition of LAB producers (e.g. *L. plantarum)* prior to a spontaneous cabbage fermentation for 10 days. Compared to standard sauerkraut fermentation conditions, aryl-lactates were enhanced by 41-125% after 10 days of fermentation with our optimization conditions (Fig S4C-D). We also found that the combination of cultures with metabolic co-factors could synergize to enhance aryl-lactate production. For example, the addition of *L. plantarum* with the aryl-pyruvate blend resulted in enhanced PLA (94%) and ILA (129%) production compared to sauerkraut control (p<0.01). Moreover, the addition of AKG and CIT had little effect on aryl-lactate production when added alone, yet these same metabolites were effective at increasing aryl-lactate levels when added alongside *L. plantarum* prior to sauerkraut fermentation (Fig S4C).

### Manipulating microbial aromatic amino acid metabolism promotes fermented food matrix capacity to activate the human aryl-hydrocarbon receptor (AhR)

We next aimed to identify the bioactivity of aryl-lactates found in fermented foods. The aryl-hydrocarbon receptor (AhR) is a ligand-dependent cytoplasmic receptor that translocates to the nucleus upon ligand binding and has been recognized as a key regulator in homeostatic processes at barrier sites, such as in the intestines^38^. Many indole compounds that are derived from the ArAA tryptophan have been identified as AhR ligands^38^. To determine which aryl-lactates activate AhR, we added increasing concentrations of aryl-lactates (0.1µM-1mM) to an *ex vivo* AhR reporter cell line (HepG2 Lucia, Invivogen), and found only ILA enhances AhR activation at concentrations ≥100 µM (p<0.05), whereas PLA and 4HPLA failed to activate the receptor (p>0.05; **Fig. 5A**).

**Figure 5.**
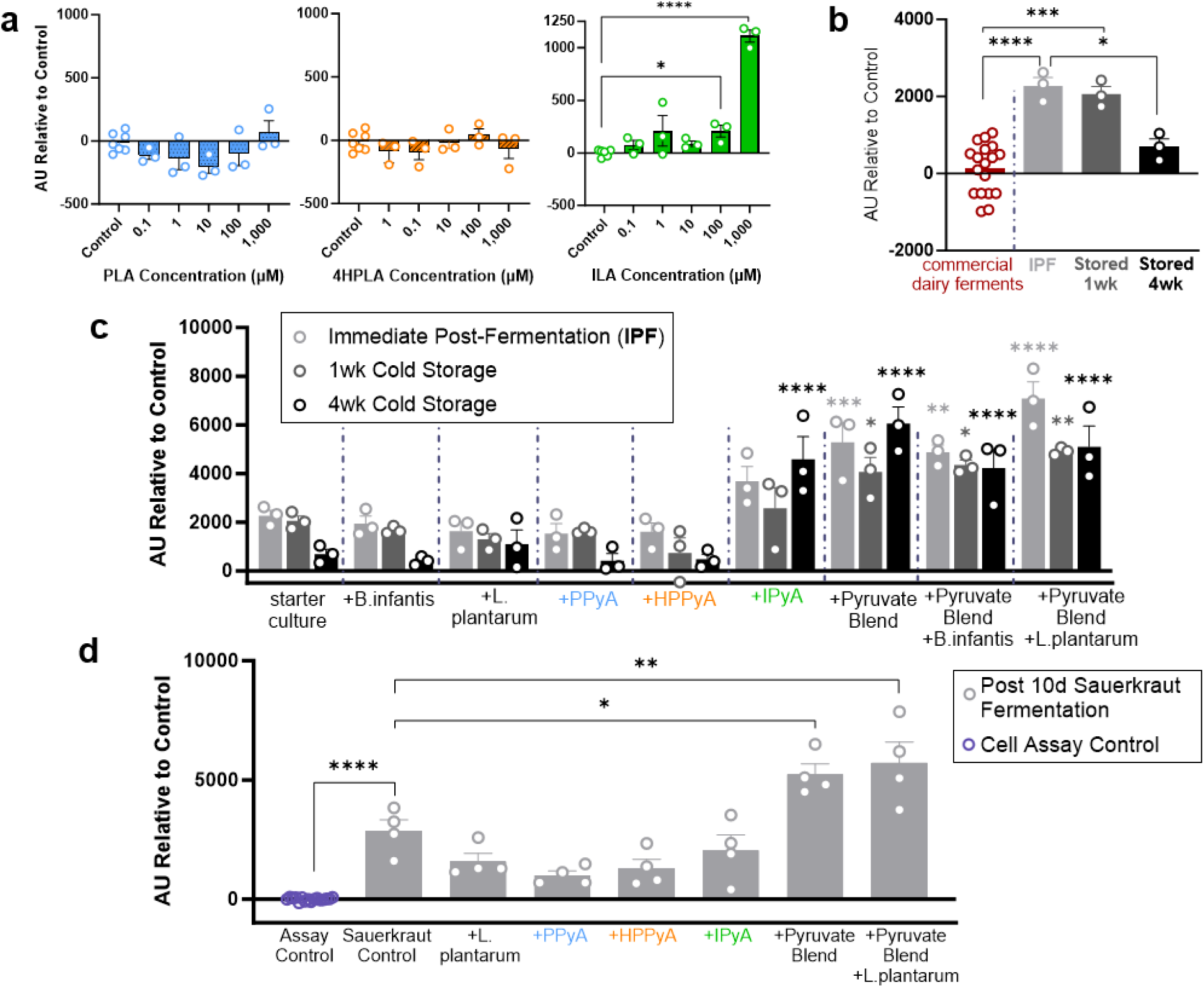
Optimized food matrix conditions to favor aryl-lactate production promotes aryl-hydrocarbon receptor (AhR) activity within two whole food matrices. (**a**) HepG2 AhR reporter cell line activity in response to aryl-lactate metabolites: phenyllactic acid (PLA), 4-hydroxyphenyllactic acid (4HPLA), and indole-3-lactic acid (ILA). Data is relative to PBS (PLA, 4HPLA) or DMSO (ILA) assay control. Statistics using 1-way ANOVA with Kruskal-Wallis post-hoc, only significant (p<0.05) p-values shown. (**b**) AhR activity in response to 6 commercially sold dairy products (left) and in-house made yogurt with starter culture only (*S. Thermophilus, L. bulgaricus*) immediately post-5hr fermentation (IPF), fermentation with 1wk cold storage, and fermentation with 4wk cold storage. Statistics using 1-way ANOVA. (**c**) AhR activity in response to optimized yogurts with various treatments and 3 timepoints: immediate post 5hr fermentation, 5hr fermentation with 1wk cold storage, and 5hr fermentation with 4wk cold storage. Treatments include starter culture only, or starter culture with additional *L. plantarum*/*B. infantis*, phenylpyruvate (PPyA), hydroxyphenylpyruvate (HPPyA), indole pyruvate (IPyA), and pyruvate blend (PPyA+HPPyA+IPyA) with and without *L. plantarum/B. infantis*. Statistical significance using 2-way ANOVA with Dunnett’s multiple comparisons test with all treatments compared to yogurt with starter culture at same timepoint showed. (**d**) AhR activity in response to sauerkraut fermentation after 10d wild ferment with various treatments: wild sauerkraut control, and additions of the following on day 0 of ferment: *L. plantarum*, PPyA, HPPyA, IPyA, and pyruvate blend with and without *L. plantarum*. Statistical significance using 1-way ANOVA comparing all treatments to sauerkraut control as well as sauerkraut control compared to cell assay control. Only significant (p<0.05) p-values shown. *p<0.05, **p<0.01, ***p<0.001, ****p<0.0001. All data displayed as mean +/- SEM.

We next investigated whether whole food matrices exhibited activity against human AhR. We first compared AhR activity in 9 store-bought dairy ferments compared to an in-house laboratory made yogurt with common starter cultures as described previously utilizing HepG2 AhR cells. With our lab-made yogurt, AhR activity was measured at three (3) time points: 1) immediate post-5hr fermentation, 2) 1 wk cold storage and 3) 4 wk cold storage. We found that AhR activity in commercial ferments was detectable, but also highly variable. Some fermented products exhibited AhR activity, while others either showed AhR inhibitory activity or had the same AhR activity as the assay control (**Fig. 5B**). Our laboratory-produced yogurt containing starter culture exhibited higher AhR activity compared to all commercial dairy ferments immediately post fermentation (**Fig. 5B**; p<0.001). However, after 4 weeks of cold storage, AhR activity declined to levels within range of commercial dairy ferments (**Fig. 5B**; p>0.05).

Next, we treated HepG2 AhR cells with yogurts optimized to favor production of aryl-lactates. Compared to the decrease in AhR activity after 4wk of storage in yogurt with starter culture only, certain yogurt treatments, including added *L. plantarum*, IPyA, and pyruvate blend with and without added *B. infantis* or *L. plantarum*, mitigated the decrease of AhR activity over time, resulting in steady AhR activity over 4wk of cold storage (**Fig. 5C**). Most notably, after 4wk of cold storage, the yogurts treated with IPyA, a metabolite we previously found to increase ILA, enhanced yogurt AhR activity more than 6-fold compared to a yogurt inoculated with starter culture only (**Fig. 5C**; p<0.0001). Furthermore, yogurt with added aryl-lactate producing bacteria (*L. plantarum*, *B. infantis*) in conjunction with added pyruvate blend significantly enhanced AhR activity immediately post fermentation period and throughout cold storage (**Fig. 5C**; p<0.001). This suggests that optimizing products to increase the production of aryl-lactates, especially ILA, will preserve biological activity of a whole food matrix over extended periods of storage.

To strengthen our observations, we next tested whether our strategies to increase aryl-lactate production could similarly enhance food AhR activity in a different food matrix, sauerkraut. Using the same AhR reporter assay described above, we first found that our control sauerkraut ferment elicited significant AhR activity as compared to media control, even without added treatments (**Fig. 5D**; p<0.0001). Encouragingly, we found that manipulated fermentation conditions that favored aryl-lactate production, most notably the addition of an aryl-pyruvate blend with an aryl-lactate producing *L. plantarum,* further enhanced AhR activity of sauerkraut compared to the control wild ferment (**Fig. 5D**; p=0.0075). Therefore, bioactivity of fermented foods, exemplified here in both yogurt and sauerkraut, can be enhanced with conditions that shift microbial ArAA metabolism towards aryl-lactate production.

## DISCUSSION

Fermented foods have grown in popularity in parallel to claims that their consumption can be health promoting, yet the compounds responsible for these effects have yet to be fully elucidated. Here, we hoped to bridge this gap by taking a closer look at microbial metabolites within fermented foods that have immunomodulatory potential. We demonstrated variations in aryl-lactate concentrations among different commercial fermented products, identified LAB that are producers of aryl-lactates, developed techniques to optimize aryl-lactate production within monoculture and whole-fermented food matrices, and confirmed biological activity of the foods optimized to have enhanced concentrations of aryl-lactates using *in vitro* methods. Ultimately, this study revealed the potential of fermented foods to act as a medium for consumption of aryl-lactates with prospects to modify human health.

Within commercial fermented foods, we observed variation of total aryl-lactate concentrations in food types (e.g., yogurt; 3.75-30 µg/mL total aryl-lactates), likely related to differences in the LAB strains present within the various brands. We found *L. plantarum* to not only be a high producer of aryl-lactates in monoculture, but also identified it as a LAB present in many of the examined commercial fermented foods, which is corroborated elsewhere^26, 27, 39–41^. *L. plantarum* has also been identified as a highly malleable LAB which likely explains its presence in a wide variety of fermented foods^42, 43^. While we also found *B. infantis* as capable of producing aryl-lactates, this bacteria species is mainly studied in the context of the microbiome and has not been investigated much in the context of fermented foods^20, 25, 34, 44^.

We found that aryl-lactate profiles varied depending on food origin – e.g., dairy versus vegetable ferments. Dairy products exhibited high levels of 4HPLA (vs. PLA), whereas vegetable ferments exhibited higher levels of PLA. This phenomenon was present in both commercial fermented foods, and the laboratory-grown sauerkraut and yogurt controls. Others have investigated presence of PLA in vegetable ferments, specifically in Chinese pickles^45^ and sauerkraut^24^, but not in relation to other aryl-lactates 4HPLA and/or ILA, making this research the first to indicate this difference. We predict that the dairy starter culture (*S. thermophilus* and *L. bulgaricus*) may have enhanced affinity to metabolize Tyr to 4HPLA in comparison to the bacteria found in plant ferments, which appear to have increased affinity to metabolize Phe to PLA^46^. Optimization techniques used herein had relatively lower impact on 4HPLA production versus other aryl-lactates in both yogurt and sauerkraut matrices (e.g., 587% increase in 4HPLA in yogurt with added *L. plantarum* + pyruvate blend immediate post-fermentation in comparison to 2,260% increase in PLA and 3,312% increase in ILA). We hypothesize this lack of sensitivity in yogurt may be related to the already robust 4HPLA producing capacity of LABs found in the food matrix, and in sauerkraut, but this should be tested more rigorously in future studies.

Similarly, we found that despite ILA levels being the lowest concentrated aryl-lactate in both commercial foods and laboratory-grown ferments, this metabolite changed the most with our optimization techniques. For example, yogurt with added IPyA exhibited 2,566% increase in ILA while yogurt with added PPyA exhibited only a 183% in PLA concentrations versus control. With this data, we predict that fLDH has preferred substrates depending on the different LABs found within food sources, with LAB within dairy preferring Tyr and HPPyA and LAB within vegetable ferments preferring Phe and PPyA. While LAB is a grouped term for select bacteria common in fermented products, it is apparent that the functional traits of these bacteria vary tremendously, ultimately influencing the use and preference of substrates within a food matrix^6^. Furthermore, these differences allow food manufacturers to tailor the production of specific aryl-lactates based on their respective bioactivity.

Microbial ArAA metabolism and aryl-pyruvates have mainly been studied in fermented foods related to taste and flavor formation^47, 48^. Herein, we demonstrate that promoting microbial ArAA metabolism to produce aryl-lactates can enhance bioactivity of food matrix as measured by human aryl-hydrocarbon receptor (AhR) activity. We also demonstrated that fermented food AhR bioactivity can be predictably preserved throughout four weeks of cold storage by enhancing microbial ArAA metabolism. This contrasts with store-bought food sources that showed highly variable AhR bioactivity, with many foods below detectable activity. Therefore, optimization of fermented foods for aryl-lactate production has the potential to improve bioactivity of foods over typical shelf life. This may have important implications for human health, as depleted microbial AhR activity has recently emerged as a key sign of chronic inflammatory diseases, including inflammatory bowel diseases and obesity^49, 50^. We thus envision that optimizing fermented foods for enhanced production of aryl-lactates to promote AhR activity may be a unique tool for combating microbial dysbiosis associated with chronic disease, but this needs to be tested.

We acknowledge the limitations of our study. While aryl-lactates are found naturally in fermented foods, we have not profiled them for potential taste modifications in whole food matrices. Furthermore, whether fermented foods optimized for enhanced production of aryl-lactates can beneficially impact human immune function and inflammatory disease states remains to be characterized.

## METHODS

### Food homogenization and metabolomics preparation

Commercially available fermented foods including kefir, yogurts, cottage cheese, sauerkraut, kimchi, pickles, fermented beets and carrots, kombucha, salami, fish sauce, and miso paste were collected from a local grocery store (brands and product descriptions listed in key resources table). All foods were diluted 1:10 with PBS and placed in a stomacher (Seward™ Stomacher™ Model 80 Biomaster Blender 110V) for 60 seconds on high and homogenized liquid was syringe filtered (0.22 -micron) before downstream LC/MS/MS metabolomics analysis.

### Liquid chromatography-mass spectrometry (LC/MS/MS)

Targeted analysis of aromatic amino acid metabolite from food samples/bacterial supernatants was performed by the Carver Metabolomics Core facility of the Roy J. Carver Biotechnology Center at UIUC. Briefly, chromatography was performed on a Vanquish UHPLC system (Thermo Scientific) with Gemini C6-Phenyl 110A column, 2 x 100 mm (3μ) column (Phenomenex); flow rate 600 μL/min. Mobile phases: 10mM Ammonium acetate (A), 0.1% formic acid in Methanol (B). Gradient: 0-0.5min – 0%B, 0.5-3min – 100%B, 3-4.1min – 100%B, 4.1-5.5min – 0%B. The injection volume was 5 μL with the column chamber temperature at 40^0^C. Mass Spectrometry was conducted with a TSQ Altis MS/MS system (Thermo Scientific). Data was acquired in positive and negative SRM modes with peak integration and quantitation using Thermo TraceFinder software.

### Real-time PCR

Store-bought fermented foods were prepared as described above. Using a spiral plater, MRS agar plates were streaked with diluted fermented food filtrates and cultured in an anaerobic chamber for 24-48 hours. Bacteria isolates were aseptically transferred to liquid MRS and grown in an incubator for an additional 24-48 hours. Bacteria isolates then underwent DNA isolation using AllPrep Bacterial DNA/RNA/Protein Kit (Qiagen) according to manufacturer’s instructions. Rt-PCR was completed with Power SYBR Green Master Mix (Thermo Fisher Scientific) and Lactobacillus primers from Kim et al^32^. Differences in gene expression were determined by Real-Time PCR (Quantstudio 5, Thermo Fisher Scientific). Raw cycle threshold (CT) values determined abundance of relative Lactobacillus species.

### Bacterial culture

Twenty strains (*Leucnonostoc citreum, Leuconostoc mesenteroides, Wiseella confusa, Lactococcus lactis, Streptococcus thermophilus, Bifidobacterium infantis, Lactobacillus* species *bulgaricus*, *plantarum, rhamnosus, and gasseri*) were evaluated for aryl-lactate producing capacity. Lactobacilli and bifidobacteria were inoculated in deMan, Rogosa and Sharp (MRS) broth while streptococci were cultured in M17 broth supplemented with 10% lactose. The cultures were incubated at 37°C or 30°C for 24 hours and 0.1% was transferred to 10 ml of sterile media and further incubated for 17 hours at same temperatures. Following incubation, samples were centrifuged at 16,102xg for 6 minutes, and cell-free supernatant was collected and filtered through 0.22μm membrane. All samples were stored at a -20°C until they were submitted to the metabolomics core for targeted liquid chromatography-mass spectrometry analysis (LC/MS/MS). Two strains (*Lactobacillus plantarum* ATCC 14917 and *Bifidobacterium infantis* ATCC 15697) were cultured as previously described. Subsequently, we evaluated the ability of these strains to produce aryl-lactates in the presence of specific key cofactors (5 mM): alpha-ketoglutarate (AKG), trisodium citrate dehydrate (CIT), and aryl-pyruvates [phenylpyruvic acid (PPyA), 4-hydroxyphenylpyruvic acid (HPPyA), and indole-3-pyruvic acid (IPyA)]. Samples were taken as previously described after 24 hours of growth in the supplemented media.

### Yogurt Preparation

Yogurt was produced in 1.5 ml centrifuge tubes to test aryl-lactate optimization by the addition of cofactors (AKG, CIT, PPyA, HPPyA, and IPyA) alone or combined with strains ATCC 14917 and ATCC 15697. Yogurt samples were prepared by reconstituting a 15% w/v mixture of skim milk powder (Millipore Sigma Skim Milk) with deionized water. The mixture first went through a heat treatment of 85°C for 30 minutes. Subsequently, the milk temperature was cooled down to 40°C before adding 0.44% starter culture (YF-L706 starter culture; *Lactobacillus delbrueckii* subsp*. bulgaricus, Streptococcus thermophilus)*. Following inoculation, cofactors were added individually, in combination, or together with 2.5% each fresh overnight cultures v/v strains ATCC 14917 and ATCC 15697 before fermentation occurred at 42°C for 5 hours. After fermentation and various time points (1- and 4-week storage at 4°C), samples were centrifuged at 16,102xg for 6 minutes, and the resulting acid whey was collected by filtration through a 0.22μm membrane. Samples were stored at -20°C until submission for LC/MS/MS.

### Sauerkraut Preparation

Sauerkraut was produced in 50 mL Falcon tubes to test aryl-lactate optimization by the addition of cofactors (AKG, CIT, PPyA, HPPyA, IPyA) alone or combined with strains ATCC14917 and ATCC15697. Each cabbage was thoroughly washed, and its outer leaves were discarded. The cleaned cabbage was finely sliced, and salt (2.5% w/w) was evenly distributed over slices. After allowing them to rest for one hour, brine was separated from cabbage. To ensure proper fermentation, 10 mL of collected brine was reintroduced, completely submerging the cabbage.

The cofactors were added into the samples in various combinations, either individually or concurrently with 2.5% v/v of aforementioned strains. All samples were thoroughly mixed, and solid phase microextraction (SPME) vial was inserted to ensure cabbage remained submerged during 10d fermentation period at 20°C. After fermentation period, samples were centrifuged at 16,102xg for 6 minutes, and resulting acid was filtered through 0.22μm syringe filters. Samples were stored at -20°C until metabolomics analysis.

### AhR Reporter Cell Line Assays

Human HepG2 liver carcinoma AhR-Lucia reporter cells (InvivoGen) were cultured according to manufacturer’s instructions. Aryl-lactates (PLA, 4HPLA, ILA) were reconstituted into solution with PBS (PLA, 4HPLA) or 1% DMSO (ILA), vortexed vigorously until dissolved, and serial diluted to desired concentrations. Fermented yogurt samples were prepared by centrifuging at 21,000xg for 10 minutes and supernatant was eluted through 0.22-micron filters. Sauerkraut samples were prepared by eluting liquid through 0.22-micron filters prior to being diluted 1:4 with PBS. Cell assay was performed according to manufacturer’s instructions. Briefly, cells were counted (Invitrogen Countess) and reconstituted in media at concentrations of 1.1×10^5^ cells/mL. 180 μL of reconstituted cells were treated with 20μl of sample in a 96-well plate and incubated at 37C with 5% CO_2_ for 17 hours. After incubation period, 20 μL of cell mixture was transferred to a white opaque 96-well plate and treated with 50 μL of QUANTI-Luc 4 Lucia/Gaussia (InvivoGen) and immediately read using luminescence at 0.1 second read time. Data was relativized to wells that contained cells with media alone.

### Statistical analysis

Statistical analysis was performed by SPSS version 29 (IBM). Student t-tests and one-way/two-way analysis of variance (ANOVA) were utilized to test differences in aryl-lactates or AhR activity as described in figure legends. In cases of non-normal distributions, non-parametric Kruskal-Wallis H tests were utilized, and Dunnett’s or Tukey post-hoc were implemented for multiple comparison testing. Data is expressed as mean ± SEM with statistical significance statistical alpha set *a priori* at p<0.05 for all analyses.

## Author Contributions

JA and MM conceived of experimental concepts. MK, AV and ECS contributed input to experimental design. MK, AV and ZX performed experiments, including fermented food preparations, bacterial cultures and AhR analyses. AU and MF performed metabolomics analysis and quality control. MK, AV, SD, MM and JA wrote the manuscript drafts. All authors read and approved the final manuscript.

## Acknowledgements

This study was funded by an intramural Personalized Nutrition Initiative Seed Grant from UIUC and the United States Department of Agriculture (USDA)-NIFA Grant # 2023-67017-39053 awarded to JA and MM. The funders played no role in study design, data collection, analysis and interpretation of data, or the writing of this manuscript.

## Data Availability

The raw datasets used and/or analyzed during the current study are available from the corresponding author on reasonable request.

## Key Resources

| Fermented Food Product | Brand & Product Description |
| --- | --- |
| Kefir | Lifeway Kefir – 1%, plain |
| Kefir | Lifeway Kefir – 1%, strawberry |
| Yogurt | Chobani – Greek, nonfat, plain |
| Kefir | Lifeway Kefir – 3.25%, plain |
| Yogurt | Siggi – nonfat, plain |
| Yogurt | Oatly! – plain |
| Yogurt | Yoplait – Greek, nonfat, vanilla |
| Yogurt | Siggi – whole fat, strawberry, drinkable |
| Yogurt | Yoplait – low fat, vanilla |
| Yogurt | Fage – nonfat, plain |
| Kefir | Zymosi – whole fat, passion fruit |
| Cottage Cheese | GoodCulture – whole fat |
| Cottage Cheese | Daisy – 4% |
| Yogurt | Fage – 5%, plain |
| Fermented Carrots | Zymbiotics – Ginger Zarrots |
| Sauerkraut | Claussen |
| Sauerkraut | PATCH |
| Kimchi | Wildbrine – kimchi with miso and horseradish |
| Kimchi | Zymbiotics |
| Fermented Beets | Zymbiotics |
| Sauerkraut | Bubbies |
| Sauerkraut | Zymbiotics |
| Pickles | Bubbies |
| Kimchi | Sunja's – medium spicy |
| Salami | Columbus – Italian Dry Salame |
| Fish Sauce | Red Boat |
| Kombucha | Synergy - Original |
| Miso | Miso Master – Organic Mellow White Miso |
| **Bacteria** | **Strain (if known)** *All strains from Miller Library* |
| *Lactobacillus plantarum* | MJM548 |
| *Lactobacillus plantarum* | MJM547 |
| *Lactobacillus plantarum* | MJM550 |
| *Lactobacillus plantarum* | MJM549 |
| *Lactobacillus plantarum* | ATCC 14917 |
| *Lactobacillus plantarum* | Cargill – LP-66 |
| *Lactobacillus plantarum* | ATCC 8014 |
| *Lactobacillus rhamnosus* | ATCC 53103 |
| *Leuconostoc mesenteroides* subsp. *mesenteroides* | ATCC 8293 |
| *Leucosonostoc citreum* | EFEL2700 |
| *Lactobacillus delbrueckii* subsp. *bulgaricus* | ATCC 11842 |
| *Lactococcus lactis* subsp. *lactis* | ATCC 19435 |
| *Lactococcus lactis* subsp. *cremoris* | ATCC 19257 |
| *Lactobacillus paraplantarum* | ATCC 700211 |
| *Lactobacillus delbrueckii* subsp. *delbrueckii* | ATCC 9649 |
| *Lactobacillus acidophilus* | ATCC 4356 |
| *Bifidobacterium infantis* | ATCC 15697 |
| *Lactobacillus casei* | ATCC 393 |
| *Lactobacillus paracasei* subsp. *paracasei* | ATCC 25598 |
| *Lactobacillus reuteri* | BioGaia |
| *Lactobacillus brevis* | ATCC 14869 |
| **Analytical Standards** | **Source** |
| 3-Phenyllactic Acid | Sigma-Aldrich (#92267) |
| DL-p-hydroxyphenyllactic acid | Sigma-Aldrich (#H3253) |
| DL-indole-3-lactic acid | Sigma-Aldrich (#I5508) |
| Phenylpyruvic acid | Sigma-Aldrich (#74346) |
| 4-Hydroxyphenylpyruvic acid | Sigma-Aldrich (#114286) |
| Indole-3-pyruvic acid | Sigma-Aldrich (#I7017) |
| L-Phenylalanine | Sigma-Aldrich (#PHR1100) |
| L-Tyrosine | Sigma-Aldrich (#PHR1097) |
| L-Tryptophan | Sigma-Aldrich (#51145) |
| 4-hydroxyphenylacetic acid | Sigma-Aldrich (#55206) |
| 3-indoleacetic acid | Sigma-Aldrich (#45533) |
| Indole-3-propionic acid | Sigma-Aldrich (#57400) |
| 3-phenylpropionic acid | Sigma-Aldrich (#W288918) |
| 4-hydroxyphenylpropionic acid | Sigma-Aldrich (#H52406) |
| Phenylacrylic acid | Sigma-Aldrich (#97013) |
| 4-hydroxyphenylacrylic acid | Sigma-Aldrich (#C9008) |
| 3-indoleacrylic acid | Sigma-Aldrich (#I2273) |
| **Labeled Isotopes** | **Source** |
| 4-hydroxyphenylpyruvic acid-13C9 | Sigma (#07491) |
| trans-Cinnamic acid-$\delta^2$,2,3,4,5,6-d6 | Sigma (#513962) |
| 3-indoleacetic acid-d5 chemical structure | MedChemExpress (#HY-18569S) |
| DL-3-phenyllactic acid-d3 | Toronto Research Chemicals Inc. (#P335567) |
| **Metabolic Co-Factor** | **Source** |
| Sodium phenylpyruvate | Sigma-Aldrich (#P8001) |
| 4-hydroxyphenylpyruvic acid | Sigma-Aldrich (#114286) |
| Indole-3-pyruvic acid | Sigma-Aldrich (#17017) |
| α-Ketoglutaric acid | Sigma-Aldrich (#75890) |
| Citric acid, trisodium salt dehydrate | Sigma-Aldrich (#227130010) |
| **Yogurt Starter Culture** | **Source** |
| YF-L706 | CHR-HANSEN (#704988) |
| **Cell Line** | **Source** |
| HepG2-Lucia<sup>TM</sup> AhR Cells | InvivoGen (#hpgl-ahr) |

