## Supplemental Figures S1-S4 for "Microbial aromatic amino acid metabolism is modifiable in fermented food matrices to promote bioactivity"

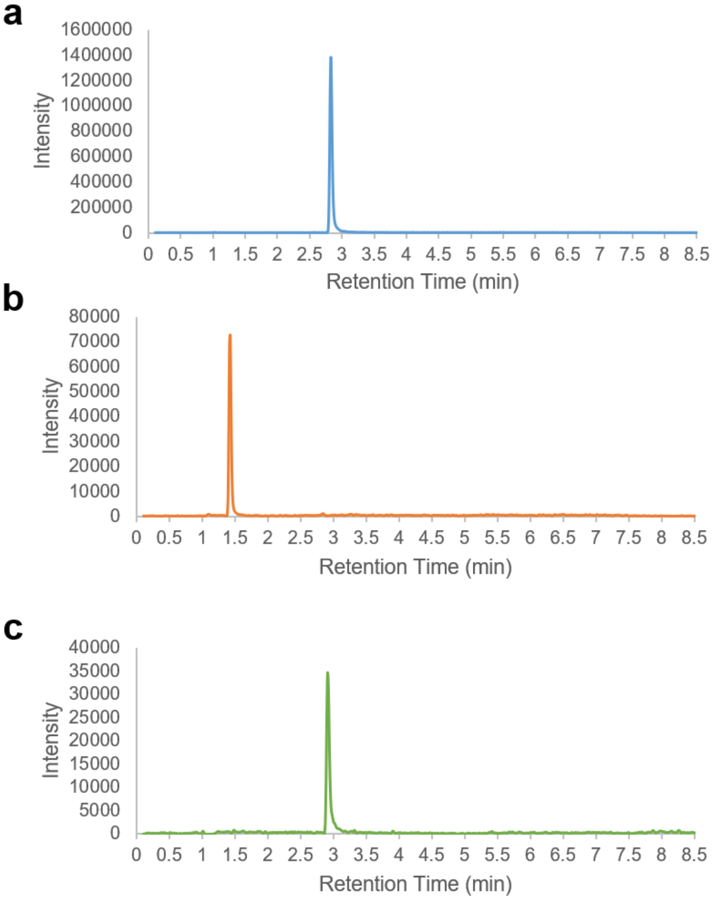


**Supplementary Figure S1** (**a-c**) chromatographs of (**a**) phenyllactic acid (PLA), (**b**) 4-hydroxyphenyllactic acid (4HPLA), and (**c**) indole-3-lactic acid (ILA) peaks of analytical standards via LC/MS/MS analysis techniques used herein.

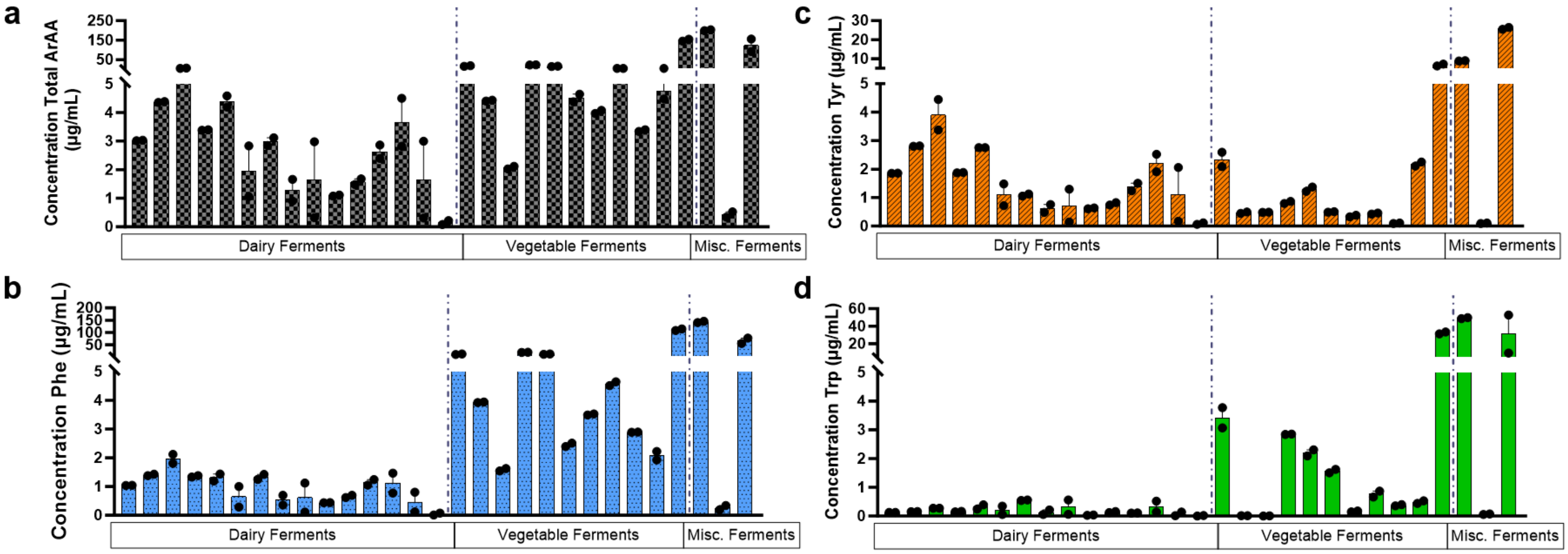


**Supplementary Figure S2**. Commercially available foods were homogenized in a stomacher for 60s, syringe filtered, and analyzed via LC/MS/MS metabolomics using aryl-lactate standards. (**a**) total aromatic amino acids (ArAAs; Phe+Tyr+Trp) (**b**) phenylalanine (Phe), (**c**) tyrosine (Tyr), and (**d**) tryptophan (Trp) concentrations are shown in µg/mL. Foods are in the same order as Fig1B-E,G, with descending total aryl-lactate concentration within the fermented food categories (dairy, vegetable, miscellaneous). A-D foods are all in the same order.

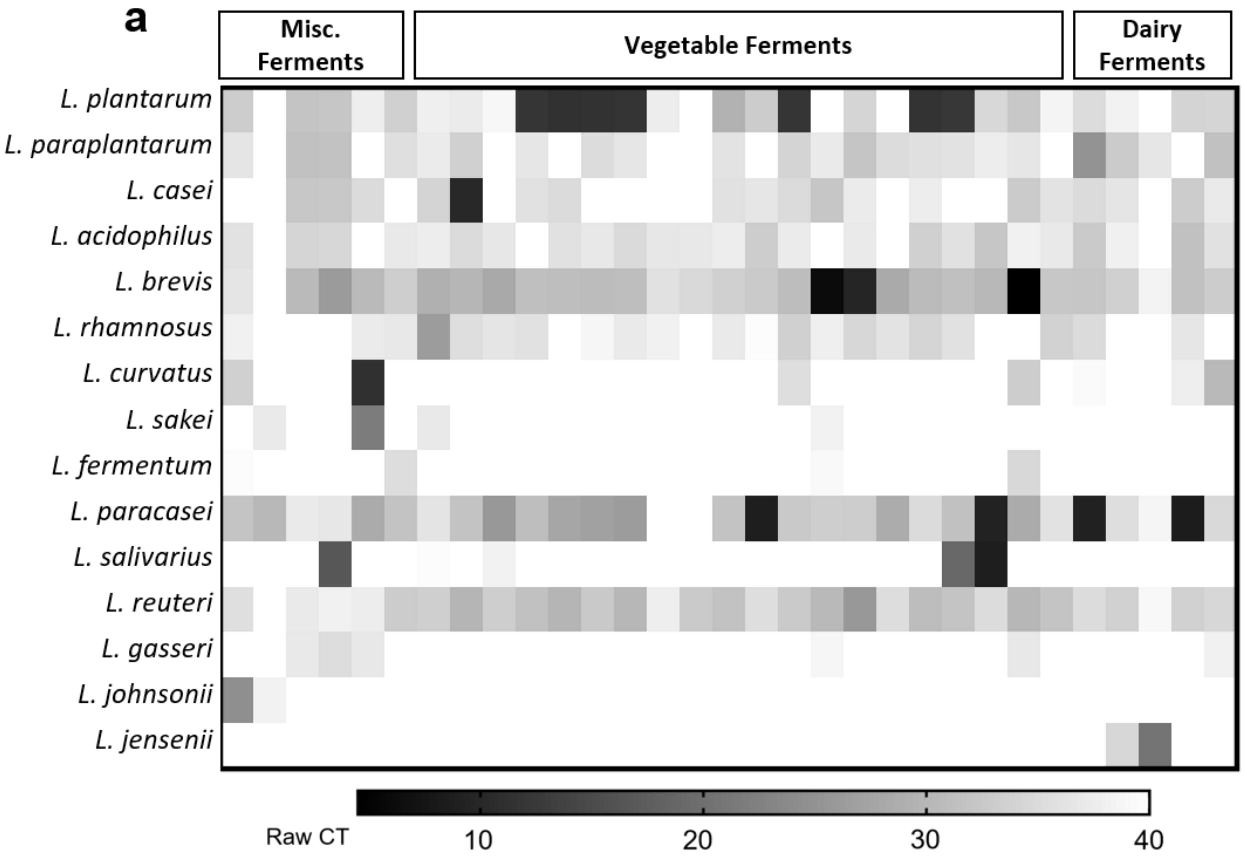


**Supplementary Figure S3**. (**a**) Commercially available fermented foods were anaerobically cultured, and isolated DNA was identified via qPCR. A lower cycle threshold (CT) indicating high abundance of the LAB species is shown in black, while a higher CT indicating low abundance or lack of the species is shown in white. All species are from the *Lactobacillus* genus.

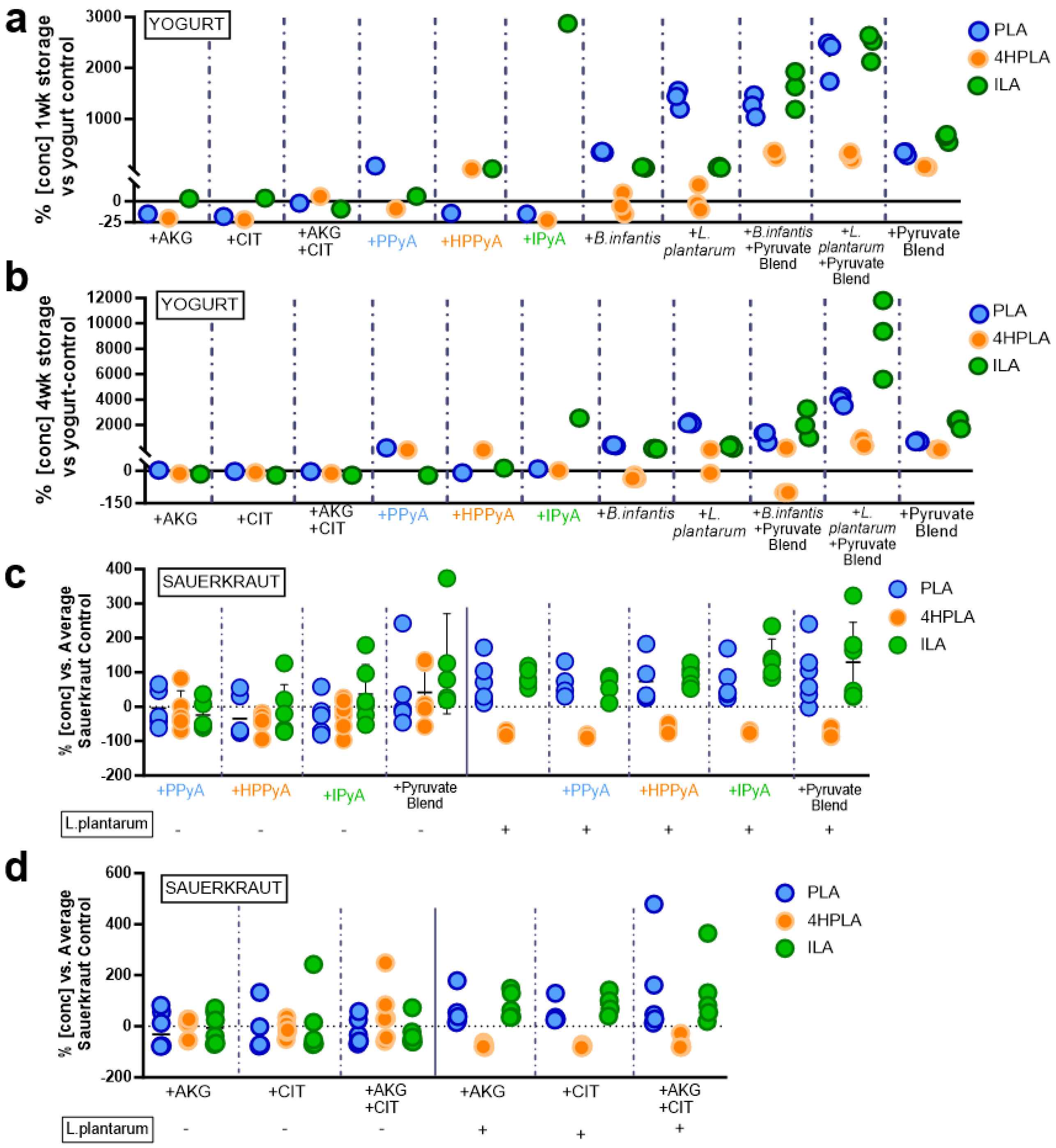


**Supplementary Figure S4**. (**a-b**) Yogurt was made by mixing milk powder and sterile water and heat treating the solution for 30min prior to being cooled to 40°C and starter culture (*Strep. Thermophilus*, *L. bulgaricus;* Chr-Hansen) plus applicable treatment were added before fermenting at 42°C for 5hr. All yogurt samples were syringe-filtered before analyzing via LC/MS/MS. Yogurt with standard starter culture was fermented for 5hr with additional (**a**) 1wk cold storage and (**b**) 4wk cold storage with treatments including: added alpha-ketoglutarate (AKG) and/or citrate (CIT), aryl-pyruvates (phenylpyruvic acid (PPyA), 4-hydroxyphenylpyruvic acid (HPPyA), indole-3-pyruvic acid (IPyA)) and *B. infantis* or *L. plantarum* with and without blend of aryl-pyruvates (**pyruvate blend**; PPyA+4HPPyA+IPyA). Phenyllactic acid (PLA), 4-hydroxyphenyllactic acid (4HPLA), and indole-3-lactic acid (ILA) percent concentration vs. yogurt control (standard starter culture only) at same timepoint is shown. (**c-d**) Sauerkraut was made by submerging freshly chopped cabbage in salt brine and fermenting for 10d. All sauerkraut samples were syringe-filtered before analyzing via LC/MS/MS. (**c-d**) Sauerkraut with no additives after a 10d ferment was compared to the following treatments: (**c**) PPyA, HPPyA, IPyA, pyruvate blend all with and without added *L. plantarum* and (**d**) AKG and/or CIT all with or without added *L. plantarum*. Data displayed as percent aryl-lactate (PLA, 4HPLA, ILA) concentration in treatment group compared to control sauerkraut. Experiment done in replicates of at least 4.
